# Determination of salivary cortisol in donkey stallions

**DOI:** 10.1101/493965

**Authors:** Francesca Bonelli, Alessandra Rota, Christine Aurich, Natascha Ille, Francesco Camillo, Duccio Panzani, Micaela Sgorbini

## Abstract

Salivary cortisol provides information about free plasma cortisol concentration and salivary sampling is a non-invasive well-tolerate procedure. The aim of this study was to validate a commercial enzyme immunoassay for the determination of salivary cortisol in donkeys.

Saliva samples were collected in 4 donkey stallions on thirteen non-consecutive days at 8:30 AM to avoid circadian variation. Animals were already accustomed to be handled. Saliva was collected by using a swab inserted at the angle of the lips, placed onto the tongue for 1 min and returned into a polypropylene tube. Tubes were centrifuged and at least 1 ml of saliva was aspirated from each sample and frozen at −20° C until analysis. A commercial enzyme immunoassay kit without extraction was used for determination of cortisol in saliva. Median cortisol concentrations with minimum and maximum value were calculated.

Recovery of cortisol standard in donkey saliva was 107.9% and serial dilution of donkey saliva samples with assay buffer resulted in changes in optical density parallel to the standard curve. Cross-reactivity of the antiserum was 10.4% with 11-deoxycortisol, 5.2% with corticosterone, 0.4% with 11-deoxycorticosterone, 0.2% with cortisone and <0.1% with testosterone, progesterone and estradiol. The intra-assay coefficient of variation was 10.7%, the inter-assay variation was 8.0% and the minimal detectable concentration was 0.01 ng/ml.

The results of the present study demonstrate the validity of a commercial kit to determine the concentration of cortisol in donkey saliva, as already reported in other species.

## Introduction

Domestic animals react to stress through physiological responses, which are the results of individual emotional reactivity. [1] The hypothalamic-pituitary-adrenocortical (HPA) system is activated in response to stressors. While early response is under catecholaminergic control, late physiological response to stressor is commonly assessed by determination of glucocorticoid concentrations, such as cortisol.

An increase in hypothalamic-pituitary-adrenocortical activity indicates a physiological response to different stressors, and measurement of plasma corticosteroids is frequently used to study middle and longtime stress response. [2]

Cortisol was primarily obtained from blood, plasma or urine. Blood sampling itself (by syringe or by catheterization) can produce stress in the animal causing a rise in cortisol levels. [3,4] Urine and fecal samples can be easily and non-invasively collected without submitting animals to stress, [5,6] but the volume/unit weight varies between animals and species [5] and time periods between application of the stressor and collection of the urine or feces cannot be easily controlled. On the other hand the animals are hardly affected by saliva sample collection, [7,8] thus saliva can be considered a non-invasive monitoring of HPA functioning in animals.

Cortisol is passively diffused into the salivary glands, not bound to proteins and its concentration is not affected by salivary flow rate. Therefore, the salivary cortisol provides direct information about free plasma cortisol concentration, which is the only biologically active fraction in the organism, owing to it being able to bind to cell receptors. [9,10]

Furthermore, the collection of saliva is a non-invasive procedure, which animals usually tolerate better than blood sampling. For all of these reasons, the interest in measurement of salivary cortisol concentration in animals has increased during the past few years.

Salivary cortisol concentration has been evaluated in human, [8,11–12] dogs, [13–16] cattle, [3,17–19] pigs [20–21] and horses. [9–10,22–25]

In the last decades other equids, the donkey, achieve a relevant position in the human society thanks to their employment in animal-assisted therapy [26] and milk production. [27–29] Therefore, the welfare and stress responses of donkey become an interesting field of research related to its behavior. To the authors’ knowledge there are no validated commercial kits for the determination of salivary cortisol levels in donkeys. We followed the hypothesis that determination of cortisol concentration in donkey saliva is possible. It was the aim of this study to validate a commercial enzyme immunoassay for the determination of salivary cortisol concentrations in donkeys.

## Materials and methods

This study was conducted between September 2015 and February 2016 and was approved by the Ethical Committee (OPBA) of the University of Pisa, according to D.lvo 26/2014. Four Amiata donkey stallions (jacks), aged 3 years (born between May and July 2012) were included in this study.

The jacks were kept together in a paddock (10 x 20 mt) for three months (September-November 2015). Then donkeys were housed for three months (December 2015-January 2016) in single boxes (3.5 x 3.5 mt) with a small outside paddock (3.5 x 6 mt), where they could see each other, but had no physical contact. Boxes were daily cleaned and bedded with straw. All the donkeys had free access to water from a water bowl and were fed meadow hay *ad libitum* and commercial concentrate (Equifioc^®^, Molitoria Val di Serchio, Lucca, Italy), according NRC recommendations for adult donkeys. [30] Weight was 280.3±30.4 kg when donkeys were moved to boxes and 283.5±27.0 kg at the end of the study, with a difference of −3, +1, +10 and +5 kg in the single animals, while Body Condition Score (BCS) was always evaluated between 5 and 6/9. [31]

Saliva samples were collected on thirteen non-consecutive days at 8:30 AM to avoid circadian variation [10] using a cotton-based swab (Salivette^®^, Sarstedt, Numbrecht-Rommelsdorf, Germany) as described in horses. [22–23] Sampling time for cortisol evaluation was scheduled in line with another project running simultaneously. [32] Briefly, all the 4 jacks underwent to semen collections thrice weekly (Tuesday, Thursday and Saturday) during 3 weeks in the month of November 2015 and February 2016. One saliva sampling took place the week before the beginning of semen collection (November 2015), while the other 12 samplings were carried out every Tuesday and Saturday when semen collection was scheduled (6 samplings in November 2015 and 6 samplings in February 2016). Thus, 7 saliva sampling were done in November 2015 and 6 in February 2017.

Samplings were always made before feeding and cleaning procedures of the boxes and before semen collection, when scheduled, with animals were at rest. Special care was used to minimize stress during the sampling procedure, in particular, the same operator handled the animals throughout the study and any special restraint during saliva samplings was applied. The swab was grasped with a surgical clamp, inserted at the angle of the lips into the mouth of the donkey and placed gently onto the tongue for 1 min and afterwards returned into a polypropylene tube. Tubes were then centrifuged for 10 min at 700 g (1800g), at least 1 ml of saliva was aspirated from each sample and the obtained saliva was frozen at −20° C until analysis.

A commercial enzyme immunoassay kit without extraction (Demeditec Diagnostics, Kiel-Wellsee, Germany) was used for determination of cortisol in saliva. The assay was validated for asinine saliva in the laboratory of the unit for Obstetrics, Gynecology and Andrology, Department for Small Animals and Horses, Vetmeduni (Vienna, Austria). Cross-reactivity of the antiserum was determined at 50% binding with 11-deoxycortisol, corticosterone, 11-deoxycorticosterone, cortisone, testosterone, progesterone and estradiol-17β. Recovery of cortisol standards added to donkey saliva was tested. Parallelism between serial dilutions of saliva samples with assay buffer and respective changes in optical density with respect to the standard curve were determined. The intra-assay and inter-assay coefficients of variation were determined from duplicates of control saliva with low and high cortisol concentrations run in each assay (n=5). The minimal detectable concentration was defined as 2 standard deviations from zero binding.

The Graph Pad Prism 6 (USA) software was used for statistical analysis. Data were analyzed for normal distribution by Kolmogorov-Smirnov test. Because not all data were normally distributed, results were expressed as median values of cortisol concentration and range calculated for each day of sampling. Values are given as ng/ml.

## Results

Saliva samples could be easily collected in all the donkeys enrolled in this study. The sampling procedure was well accepted by all the animals. In one donkey (n. 4 table 1), the amount of saliva collected was occasionally too small for analysis (samples 2nd, 6th and 9th).

**Tab 1.**
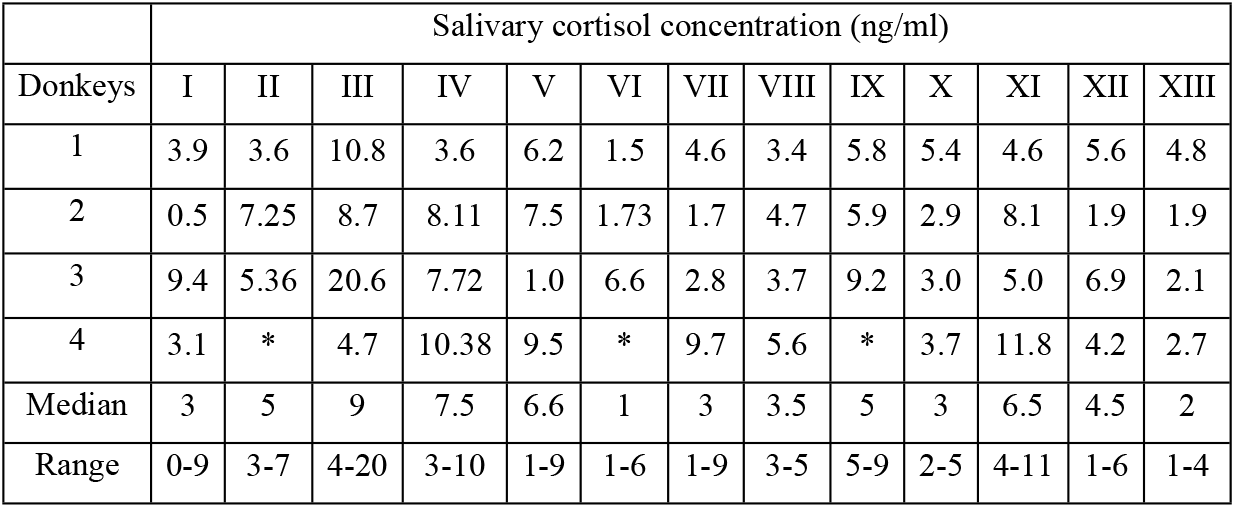
Results of salivary cortisol concentration obtained in 4 individual donkey stallions on 13 non-consecutive days. Legend: *Amount of saliva not sufficient for cortisol analysis.

Recovery of cortisol standard in donkey saliva was 107.9% and serial dilution of donkey saliva samples with assay buffer resulted in changes in optical density parallel to the standard curve. Cross-reactivity of the antiserum at 50% binding was 10.4% with 11-deoxycortisol, 5.2% with corticosterone, 0.4% with 11-deoxycorticosterone, 0.2% with cortisone and <0.1% with testosterone, progesterone and estradiol. The intra-assay coefficient of variation was 10.7%, the inter-assay variation was 8.0% and the minimal detectable concentration was 0.01 ng/ml.

Results of cortisol concentrations determined in saliva collected from donkeys on different days are presented in Table 1.

## Discussion

The production of cortisol is regulated by the corticotrophic axis and can be altered in different circumstances. Cortisol concentrations can rise in stressful situations, thus it is considered to be an indicator of animal welfare [4] Although traditionally cortisol concentrations have been determined in plasma or serum, their determination in samples easily collectable under field conditions (saliva, milk, urine, feces and hair) can be of interest for animal welfare studies in different species. [8,33–37] Thus, in our study, a commercial kit assay system, originally used in humans for measuring concentration of cortisol, was adapted and validated for donkey saliva samples.

The evaluation of salivary cortisol has some advantages in large animals. Cortisol concentration in saliva corresponds to the free fraction of cortisol in plasma, thus salivary cortisol represents a better indicator of the possible effects of the corticotrophic axis on the animal organism than plasma cortisol. [17] Moreover, blood collection always produces stress in the animal that can cause cortisol levels to rise. [7–8] On the other hand, saliva sampling, especially in already handled animals, can be an easy and stress-free procedure as confirmed by our results. All donkeys included in the study well tolerated the procedures and the amount of saliva collected was good enough in the majority of the samples. The results of the present study demonstrate the validity of a commercial kit to determine the concentration of cortisol in donkey saliva, as already reported in dogs [13–15] and horses. [9–10,22–23,38]

In the specificity tests, all the cross reactions observed were in line with the manufactory performance characteristic declared. A very low cross-reaction has been found with 11-deoxycortisol (10.4%) and corticosterone (5.2%). The 11-deoxycortisol is a direct precursor of cortisol and is found in plasma at lower concentrations than cortisol. Literature showed that the 11-deoxycortisol have none glucocorticoid effect and almost no mineral corticoid effect. [17] Corticosterone is the predominant glucocorticoid produced in some species (i.e. mice, rats and birds). [39] Even if corticosterone is also synthesized in cortisol-dominant species, like equids, its glucocorticoid activity in these species is still under assessment. [39] Cross-reactivity with 11-deoxycorticosterone, cortisone and with testosterone, progesterone and estradiol were negligible. Intra- and inter-assay coefficients of variation are used to control the potential analytic error in biological assay systems. [40] Our results showed a slightly increased intra- and inter-assay CV % compared with the manufactory performance characteristic declaration (intra-assay CV of 3.8-5.8% and inter-assay CV of 6.4-6.2%). However, the validation results indicated that the method was adequate for salivary cortisol measurements given that it is generally accepted that the CV must be lower than 20% for analytical determinations. [40] The minimal detectable concentration found in our study was slightly below those indicated by the manufactory guidelines (0.01 ng/ml *vs* 0.024 ng/ml, respectively) and published in cows, [17,41] but similar to results in pigs [20] and horses. [10]

## Conclusions

In conclusion, given the advantages of the stress-free collection and processing of saliva and the good performance of the assay, the use of cortisol detection in saliva could be considered highly suitable in donkey stress research. A limit of the present study is the fact that we did not compare salivary cortisol concentrations to concentration of cortisol in blood collected at the same time. However, when this was done in horses, the relationship was very good. [9–10] Also, despite in the majority of the samples, the quantity of saliva was good enough for running the assay, an improvement in saliva collection procedure need to be addressed.

## Acknowledgements

We are grateful to Ente Terre Regionali Toscane for allowing us to employ the animals for this study.

